# Moderate consistency in smooth newt behaviour

**DOI:** 10.1101/363648

**Authors:** Petr Chajma, Oldřich Kopecký, Jiří Vojar

**Affiliations:** Department of Ecology, Faculty of Environmental Sciences, Czech University of Life Sciences Prague, Kamýcká 129, Praha – Suchdol, 165 00, Czech Republic; Department of Zoology and Fish Farming, Faculty of Agrobiology, Food and Natural Resources, Czech University of Life Sciences Prague, Kamýcká 129, Praha – Suchdol, 165 00, Czech Republic

## Abstract

Behavioural consistency (i.e., personality) is a novel field of research in amphibians. Current published studies often address only one or two aspects of personality and therefore cannot assess more complex relationships and behavioural syndromes. This is the first study focusing on all relevant behavioural traits and their relationships in urodele amphibians. Based on the three trials of the experiment, we examined the consistency of activity (time spent moving), boldness (latency of the first movement and time spent escaping) and exploration (number of visited segments of testing arena) of 42 smooth newts (*Lissotriton vulgaris*). Individual consistency, calculated through the intraclass correlation coefficient (ICC), was low in newt activity (ICC = 0.192) and moderate in boldness (0.476) and exploration (0.403). Activity was moderately consistent for each trial (0.425), indicating a possible habituation, supported by a decrease of mean activity throughout the trials. Correlation of the behavioural traits studied suggests the presence of a behavioural syndrome, which potentially shaped the traits together. Our findings suggest the need for a complex approach to the study of amphibian personality and the need for standardized methodology, which would solve the current difficulties in comparing published results.

## Introduction

Behavioural consistency (i.e., personality) is a well-known phenomenon studied in many taxa (Gosling 2001; Sih et al. 2004; Réale et al. 2007; Garamszegi et al. 2013) and recently studied in amphibians (Aragón 2011; Koprivnikar et al. 2011; Maes et al. 2012; Wilson and Krause 2012; Brodin et al. 2013; Carlson and Langkilde 2013, 2014a). Consistency in the expression of behavioural traits over time and in different situations, as well as the correlation of those traits, i.e., behavioural syndrome (Sih et al. 2004), is often linked to survival in predator-prey situations (Dingemanse and de Goede 2004; Carlson and Langkilde 2014b) (but see Carlson and Langkilde 2014a), reproductive success (Dingemanse and Réale 2005; Cole and Quinn 2014), disease risk (Koprivnikar et al. 2011) and dispersal tendencies (Cote et al. 2010, 2013; Maes et al. 2012; Brodin et al. 2013). Therefore, animal personality plays an important part in individual life histories and should be inspected and carefully considered when dealing with most aspects of animal ecology.

Amphibian personality research, however, is limited and includes mostly studies performed on anurans (for a thorough review, see Kelleher et al. 2017). In contrast, there are considerably fewer studies for urodeles (Sih et al. 2003; Aragón 2011; Gifford et al. 2014; Winandy and Denoël 2015) and none for caecilians. Existing amphibian studies most commonly address the consistency of activity, boldness and exploration and have yet to consider the consistency of remaining personality traits – aggressiveness and sociability (Kelleher et al. 2017). Most of these studies (none of which consider urodeles), however, address only one or two of these behavioural traits (axis of personality) and usually cover some specific problem, not personality *per se*. There are also differences in approaches to behavioural consistency, because some studies cover differences across time (e.g. Wilson and Krause 2012; Maes et al. 2012; Carlson and Langkilde 2013; Brodin et al. 2013), while others consider differences across situations (e.g. Sih et al. 2003; Aragón 2011; Koprivnikar et al. 2011).

Therefore, the aim of our study was to measure the consistency of main behavioural traits—activity, exploration and boldness—in one experiment to examine the main types of behavioural responses and to focus only on temporal consistency while reducing other factors. Furthermore, we wanted to assess the correlations between these behaviour types (i.e., the existence of behavioural syndrome).

## Methods

### Experimental design

The experiment was carried out in laboratory conditions at the Czech University of Life Sciences in Prague. For a model organism, we chose the urodele that was most abundant locally, the smooth newt (*Lissotriton vulgaris*). At the start of the reproductive season in the beginning of May 2017, 21 males and 21 females were captured by nets in a single pond in the Stará Lysá village in the Central Bohemia region. The newts were housed separately in plastic containers of dimensions 18 × 12 × 14 cm that were filled with aged tap water, and the newts were fed *Daphnia* and Chironomidae larvae *ad libitum*. The air temperature in the laboratory was constant and set to 17°C. Sufficient light intensity in a diurnal cycle was provided by the translucent roof of the laboratory.

The experiment itself was conducted between 13^th^ and 27^th^ May in two experimental arenas made of non-transparent round green water barrels with bottom diameters of 80 cm. Using a non-toxic waterproof marker, a square grid of 7 cm segments was drawn at the bottom to better assess the position of each newt. The arena was filled with 5 cm of cold tap water (10.8–11.2°C), and after each recording, the water was changed, and the arena was thoroughly cleaned with a clean sponge, pressurized water and then left to dry to eliminate any potential chemical cues that remained from the previous individual tested.

Each trial of the experiment was 12 minutes long. Behaviour was recorded at 25 frames per second with a full HD camera, positioned approximately 150 cm above the water level. Newts were separately inserted under the transparent glass dome (10 cm diameter) into the centre of the arena and left to calm down for the first two minutes. Then, the dome was carefully removed in a motion perpendicular to the ground, and the recording was initiated. To measure the temporal repeatability of the behaviour, each individual was recorded three times with a six day gap between each recording, which was the longest gap possible before the newts started to shift to the terrestrial phase of the season. Unfortunately, three videos were lost due to technical difficulties in the last trial of the experiment, so the total number of analysed videos was 123.

Three types of behaviour (personality traits) were tracked: activity, exploration and boldness. Activity was measured as the amount of time [s] during which the individual moved. Furthermore, the movement activity was divided to walking and swimming to distinguish the role of each in total activity and the consistency of each as well as to determine the consistency of the choice of locomotion (i.e. the proportion of activity spent by walking). Exploration was recorded as the number of grid blocks that an individual entered. For the sake of better comparison with the other studies, boldness/shyness was measured as the latency of the first movement [s] (the most common but imprecise measure of boldness, see Discussion) as well as the time [s] spent at the outermost edge of the arena (our preferred measure). Staying in its vicinity (thigmotaxis) can be interpreted as an escape response and therefore can be a valuable measure of shyness (Burns 2008; Harris et al. 2009; Carlson and Langkilde 2013). Behaviour was scored manually by the same person using the software Observer XT v. 10 (Noldus 2010). The study was carried out in accordance with permit SZ-092744/2012KUSK/3 issued by the Regional Office of the Central Bohemian Region of the Czech Republic and approved by an institutional committee based on the institutional accreditation No. 63479/2016-MZE-17214 of Ministry of Agriculture of the Czech Republic.

### Data analysis

To test the differences in activity, time spent walking, swimming, number of visited squares (exploration), latency of the first movement (boldness) and time spent near the outermost edge of arena (shyness) between trials and sexes (independent variables), we created separate linear mixed effects models (LMM) for each mentioned characteristic (dependent variable) fitted by restricted maximum likelihood (REML) with the individual (1–42) as random intercept. Apart from mentioned variables, we also tested the dependency of the proportion of walking activity (i.e. time spent walking divided by time spent active) of each newt on the same fixed (trial and sex) and random effects (individual). This was done to assess if the preferred type of locomotion differed between sexes and trials. Albeit modelling proportions, this particular model reasonably met the assumptions for LMM.

Each model was also tested for the effect of time of day, when the experimental trial took place. Because the dependency on time is rarely linear, we decomposed this variable to sine and cosine of time in radians to accommodate for its periodical nature. Upon meeting all underlying assumptions, models were evaluated using Type II Wald Chi-squared tests. The time of day, nor the sex of the newts didn’t affect any of tested variables (see Supplementary information for the details) and were therefore not included in repeatability analyses.

Individual consistency (repeatability) in measured traits (dependent variables from previous models) was calculated using the intraclass correlation coefficient (ICC), computed from the variance components of models similar to previous ones, but with no fixed effects and the trial number as a second random variable. Note that adding trial number as a random intercept allowed us to estimate its’ consistency, i.e. the between subject similarity in measured traits expression during each trial of the experiment and estimate individual repeatability stripped from the effect of trial order.

Confidence intervals (CI) for the ICC were estimated by parametric bootstrapping with 1000 iterations (for details see Nakagawa and Schielzeth 2010). Confidence interval for the proportion of walking activity was estimated from the fixed intercept of the linear mixed effects model with the individual and trial as random intercepts, using the profile likelihood method.

The existence of behavioural syndromes was tested using Kendall’s coefficient of concordance, rather than ICC, because the interest lays in the ranks of responses, rather than their absolute value. Pairwise similarities were analysed using Pearson’s correlation coefficient. All statistical analyses were performed in R 3.3.1 (R Core Team 2016) using *lme4* (Bates et al. 2015), *car* (Fox and Weisberg 2011) and *rptR* (Stoffel et al. 2017) packages at the level of significance *α* = 0.05.

### Data Availability

The datasets analysed during the current study are available in the Open Science Framework repository, https://osf.io/nbfk6/.

## Results

### Activity

Mean activity significantly differed between trials (for details, see Supplementary information). The initial mean activity of 309.9 seconds decreased by 32 % in the second trial and by 20 % in the third trial. Although with a low ICC (0.192), individual activity was significantly repeatable. Activity among the newts was also significantly repeatable within trials of the experiment (*ICC* = 0.416) (for details, see Table 1a and Table 1b).

**Table 1a.**
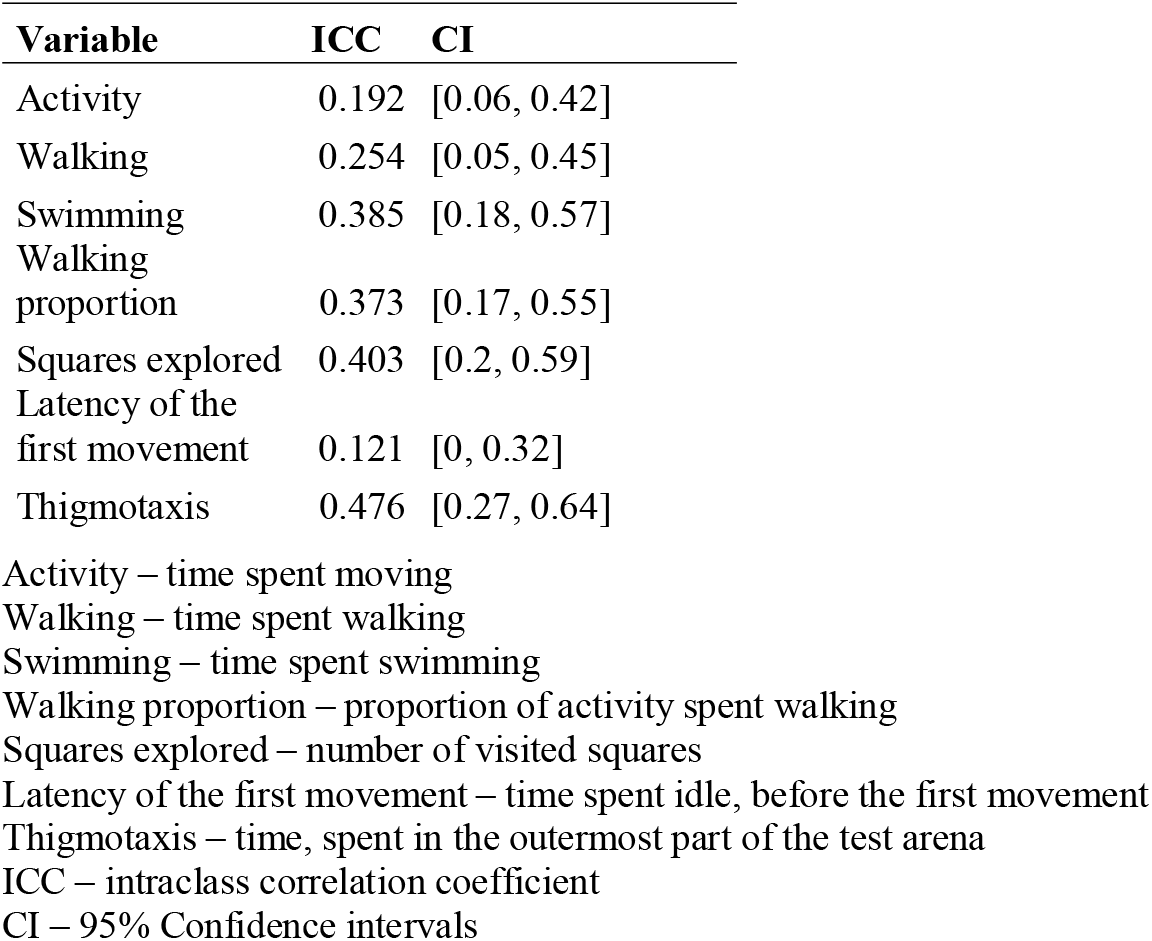
Individual repeatability of behavioural traits.

**Table 1b.**
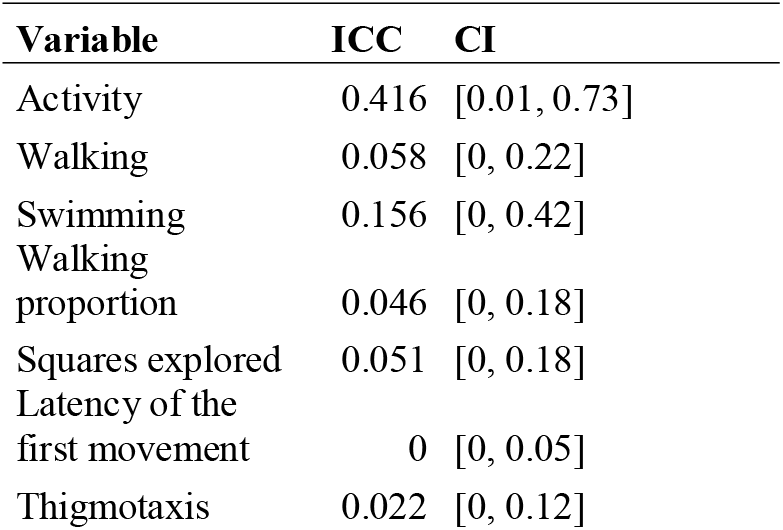
Trial repeatability of behavioural traits.

When the activity was divided into walking and swimming, walking was the preferred type of locomotion taking up 71 % of time spent active (*CI* = [0.583, 0.840]). Both the amount of walking (*P* < 0.01) and swimming (P < 10^−4^) differed significantly between the trials of the experiment. The portion of activity spent by walking, however, increased only slightly during the second and third trials (P = 0.05). Albeit significant, the repeatability of walking was relatively low for each newt (*ICC* = 0.254) and non-significant for the trial (*ICC* = 0.058). Swimming was more repeatable than walking and was significant for each individual (*ICC* = 0.385) but not each trial (*ICC* = 0.156). The proportion of walking activity was fairly consistent for individuals (*ICC = 0.373*) but not for each trial (*ICC = 0.046*) (for details, see Table 1a and Table 1b).

### Exploration

Similar to activity, there was a significant difference between each trial of the experiment (P = 0.01, see Supplementary information). The initial mean of 27.8 explored squares decreased by 2.5 % in the second and by 18 % in the third trial. Exploration was significantly repeatable for each newt with a moderate ICC (0.403). In contrast to activity, there was no repeatability for the exploration in each trial (*ICC* = 0.051, see Table 1a and Table 1b).

### Boldness

Boldness was measured as the latency to move and the time spent with an escape response. Latency to move was not dependent on any tested variables (see Supplementary information). Time spent with an escape response was marginally independent of the trial (P = 0.06). The initial mean time spent with the escape response of 226.9 seconds decreased by 5.5 % in the second trial and then rose by 30 % in the third trial. The repeatability of latency and time spent escaping was similar for each trial of the experiment, but not for individual newts. Latency was not repeatable for both newt (*ICC* = 0.121) and trial (*ICC* = 0). Time spent escaping was, on the other hand, moderately repeatable for each newt (*ICC* = 0.476) and not repeatable for each trial (*ICC* = 0.022, see Table 1a and Table 1b).

### Correlated behaviour

The similarity in mean activity, exploration and time spent escaping (shyness) of individuals was relatively high (*Kendall’s W* = 0.716, *P* < 10^−13^). Pairwise correlations showed a strong positive relationship between activity and time spent escaping (*r* = 0.734, *P* < 10^−6^) and a moderate correlation between activity and exploration (*r* = 0.539, *P* = 0.0002) and between time spent escaping and exploration (*r* = 0.405, *P* = 0.0078).

## Discussion

Observed behavioural responses of studied newts were moderately individually consistent for swimming activity, proportion of walking activity, exploration and escape response (thigmotaxis). Repeatability of activity as a whole was lower because of the less repeatable walking activity that was more prevalent than swimming activity. The consistency of the proportion of walking, i.e., the choice of locomotion type, however, suggests that even when activity levels change, the preferred type of locomotion does not. The activity was moderately consistent for each trial, meaning it decreased for all newts similarly during each trial of the experiment. Boldness was consistent only if measured as thigmotaxis (i.e., time spent with escape response), not as the latency of the first movement. Studied behaviour responses also did not differ between sexes (as opposed to Aragón 2011) and were unaffected by the time of day when experiment started.

According to recent review of Kelleher et al. (2018), a total of six studies tested the consistency of expressed behavioural traits in larval and eight in post-metamorphic amphibians. Up to date, there is, however, no review comparing the magnitude of behavioural trait consistency of said studies. Therefore, in attempt to compare our results to the remaining works, we reviewed available literature in Table 2. Out of 16 reviewed studies, five addressed urodeles, but only three provided repeatability estimates, none of which had more than two repeated measurements. Furthermore only one of described urodele studies provided evidence for temporal repeatability of a personality trait (exploration) (Gifford et al. 2014). If we include the works that compare behaviour in different odour treatments (no, conspecific or predator odour), observed repeatability in our study was lower for general activity and similar for boldness and exploration, though the unbiased comparison is impossible, due to substantial differences in used methods.

**Table 2.**
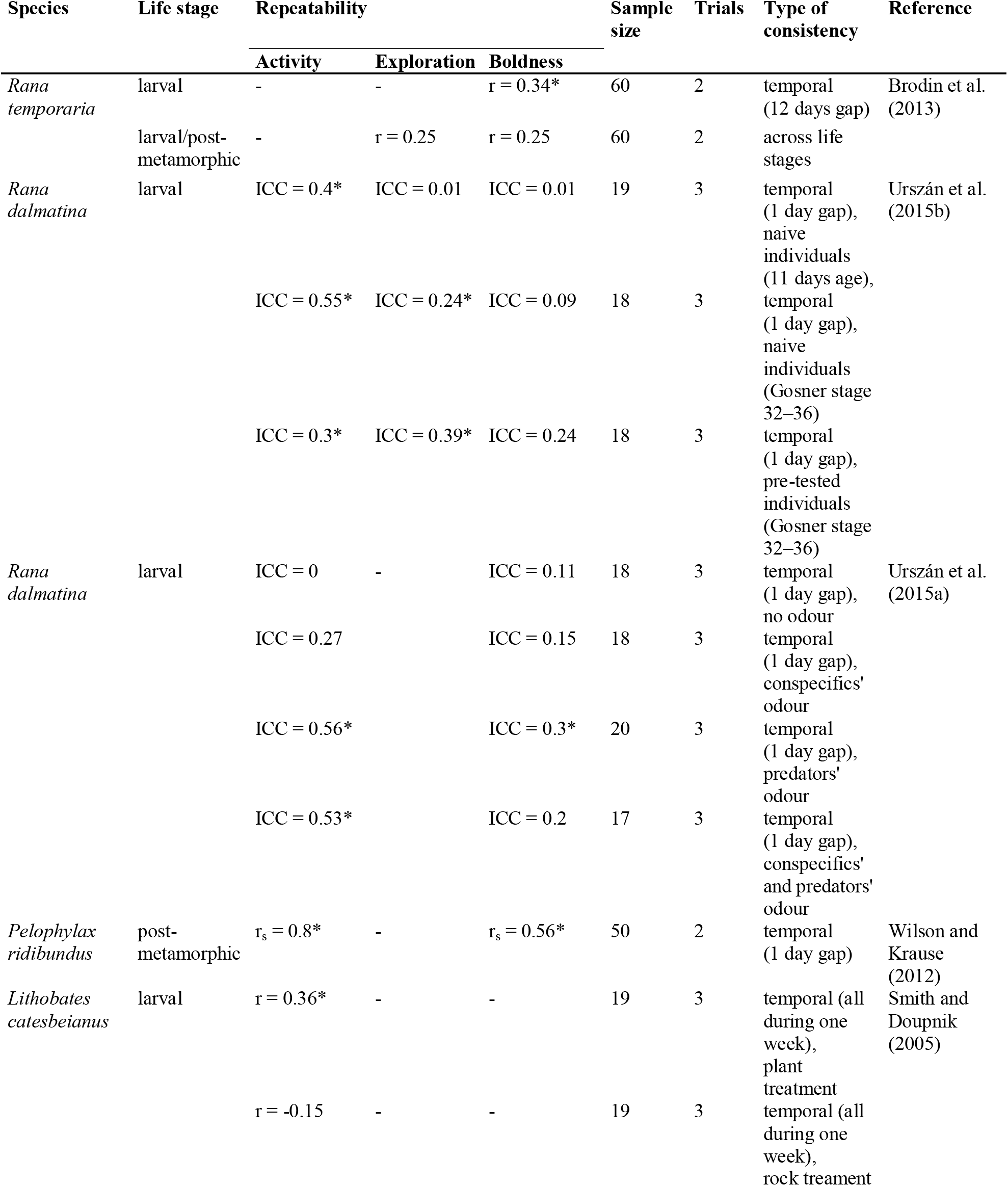
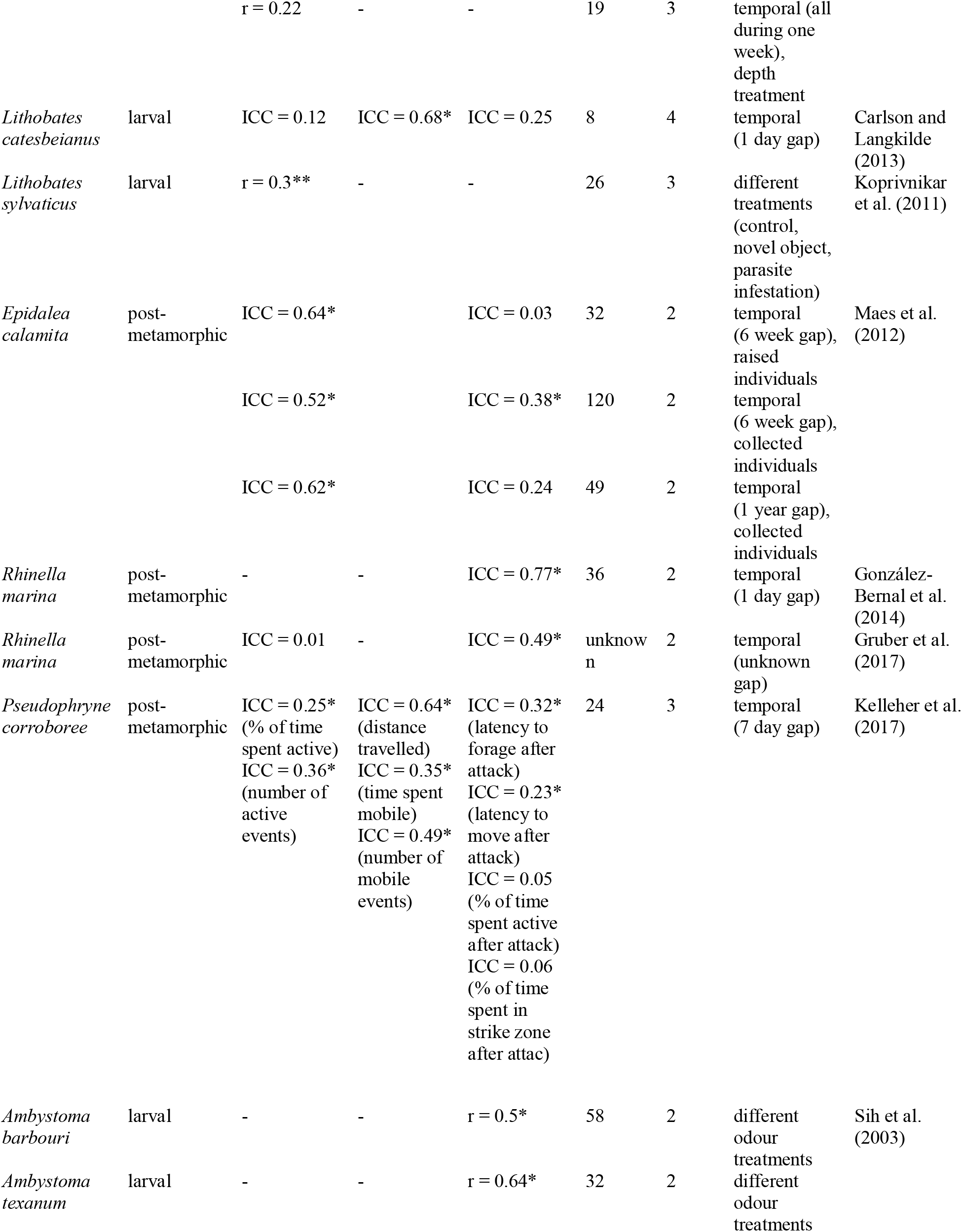
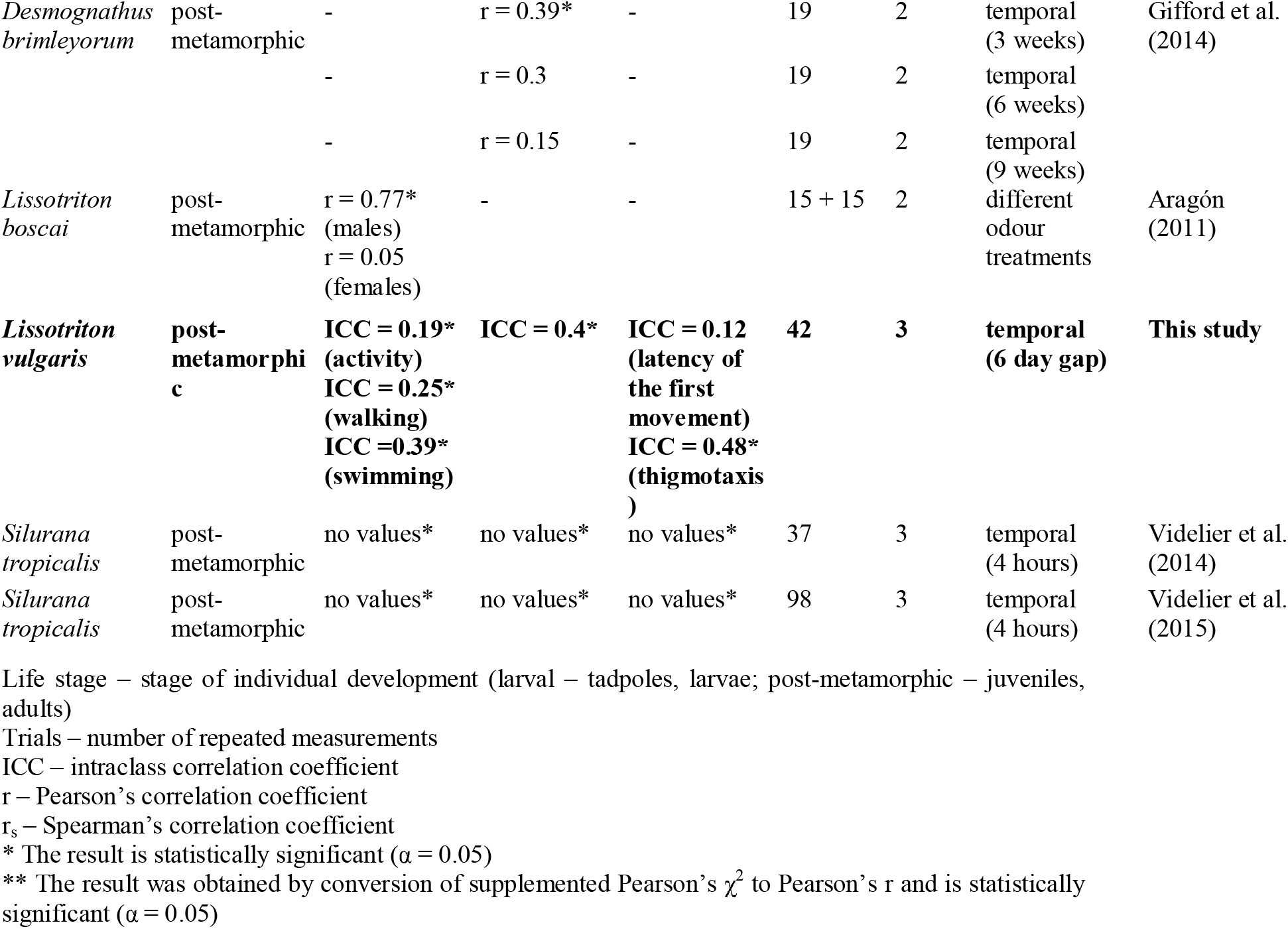
Summary of amphibian personality studies effect sizes.

| Species | Life stage | Repeatability |  |  | Sample size | Trials | Type of consistency | Reference |
| --- | --- | --- | --- | --- | --- | --- | --- | --- |
|  |  | Activity | Exploration | Boldness |  |  |  |  |
| *Rana temporaria* | larval | - | - | $r = 0.34^*$ | 60 | 2 | temporal (12 days gap) | Brodin et al. (2013) |
| | larval/post-metamorphic | - | $r = 0.25$ | $r = 0.25$ | 60 | 2 | across life stages | |
| *Rana dalmatina* | larval | ICC = 0.4* | ICC = 0.01 | ICC = 0.01 | 19 | 3 | temporal (1 day gap), naive individuals (11 days age), | Urszán et al. (2015b) |
|  |  | ICC = 0.55* | ICC = 0.24* | ICC = 0.09 | 18 | 3 | temporal (1 day gap), naive individuals (Gosner stage 32–36) |  |
|  |  | ICC = 0.3* | ICC = 0.39* | ICC = 0.24 | 18 | 3 | temporal (1 day gap), pre-tested individuals (Gosner stage 32–36) |  |
| *Rana dalmatina* | larval | ICC = 0 | - | ICC = 0.11 | 18 | 3 | temporal (1 day gap), no odour | Urszán et al. (2015a) |
|  |  | ICC = 0.27 |  | ICC = 0.15 | 18 | 3 | temporal (1 day gap), conspecifics' odour |  |
|  |  | ICC = 0.56* |  | ICC = 0.3* | 20 | 3 | temporal (1 day gap), predators' odour |  |
|  |  | ICC = 0.53* |  | ICC = 0.2 | 17 | 3 | temporal (1 day gap), conspecifics' and predators' odour |  |
| *Pelophylax ridibundus* | post-metamorphic | $r_s = 0.8^*$ | - | $r_s = 0.56^*$ | 50 | 2 | temporal (1 day gap) | Wilson and Krause (2012) |
| *Lithobates catesbeianus* | larval | $r = 0.36^*$ | - | - | 19 | 3 | temporal (all during one week), plant treatment | Smith and Doupnik (2005) |
| | | $r = -0.15$ | - | - | 19 | 3 | temporal (all during one week), rock treatment | |
| | | $r = 0.22$ | - | - | 19 | 3 | temporal (all during one week), depth treatment | |
| *Lithobates catesbeianus* | larval | ICC = 0.12 | ICC = 0.68* | ICC = 0.25 | 8 | 4 | temporal (1 day gap) | Carlson and Langkilde (2013) |
| *Lithobates sylvaticus* | larval | $r = 0.3^{**}$ | - | - | 26 | 3 | different treatments (control, novel object, parasite infestation) | Koprivnikar et al. (2011) |
| *Epidalea calamita* | post-metamorphic | ICC = 0.64* |  | ICC = 0.03 | 32 | 2 | temporal (6 week gap), raised individuals | Maes et al. (2012) |
|  |  | ICC = 0.52* |  | ICC = 0.38* | 120 | 2 | temporal (6 week gap), collected individuals |  |
|  |  | ICC = 0.62* |  | ICC = 0.24 | 49 | 2 | temporal (1 year gap), collected individuals |  |
| *Rhinella marina* | post-metamorphic | - | - | ICC = 0.77* | 36 | 2 | temporal (1 day gap) | González-Bernal et al. (2014) |
| *Rhinella marina* | post-metamorphic | ICC = 0.01 | - | ICC = 0.49* | unknown | 2 | temporal (unknown gap) | Gruber et al. (2017) |
| *Pseudophryne corroboree* | post-metamorphic | ICC = 0.25* (% of time spent active)<br>ICC = 0.36* (number of active events) | ICC = 0.64* (distance travelled)<br>ICC = 0.35* (time spent mobile)<br>ICC = 0.49* (number of mobile events) | ICC = 0.32* (latency to forage after attack)<br>ICC = 0.23* (latency to move after attack)<br>ICC = 0.05 (% of time spent active after attack)<br>ICC = 0.06 (% of time spent in strike zone after attack) | 24 | 3 | temporal (7 day gap) | Kelleher et al. (2017) |
| *Ambystoma barbouri* | larval | - | - | $r = 0.5^*$ | 58 | 2 | different odour treatments | Sih et al. (2003) |
| *Ambystoma texanum* | larval | - | - | $r = 0.64^*$ | 32 | 2 | different odour treatments | |
| *Desmognathus brimleyorum* | post-metamorphic | - | r = 0.39* | - | 19 | 2 | temporal (3 weeks) | Gifford et al. (2014) |
|  |  | - | r = 0.3 | - | 19 | 2 | temporal (6 weeks) |  |
|  |  | - | r = 0.15 | - | 19 | 2 | temporal (9 weeks) |  |
| *Lissotriton boscai* | post-metamorphic | r = 0.77* (males)<br>r = 0.05 (females) | - | - | 15 + 15 | 2 | different odour treatments | Aragón (2011) |
| *Lissotriton vulgaris* | post-metamorphic | ICC = 0.19* (activity)<br>ICC = 0.25* (walking)<br>ICC = 0.39* (swimming) | ICC = 0.4* | ICC = 0.12 (latency of the first movement)<br>ICC = 0.48* (thigmotaxis) | 42 | 3 | temporal (6 day gap) | This study |
| *Silurana tropicalis* | post-metamorphic | no values* | no values* | no values* | 37 | 3 | temporal (4 hours) | Videliér et al. (2014) |
| *Silurana tropicalis* | post-metamorphic | no values* | no values* | no values* | 98 | 3 | temporal (4 hours) | Videliér et al. (2015) |

Life stage – stage of individual development (larval – tadpoles, larvae; post-metamorphic – juveniles, adults)
Trials – number of repeated measurements ICC – intraclass correlation coefficient r – Pearson's correlation coefficient $r_s$ – Spearman's correlation coefficient \* The result is statistically significant ( $\alpha = 0.05$ ) \*\* The result was obtained by conversion of supplemented Pearson's $\chi^2$ to Pearson's r and is statistically significant ( $\alpha = 0.05$ )

### Activity

Taking into account all of the reviewed studies for both anuran and urodelan amphibians, the repeatability of activity varied odour treatments (Urszán et al. 2015a), arena structure (Smith and Doupnik 2005), breeding origin – wild or captive (Maes et al. 2012), sex (Aragón 2011), previous experience (Urszán et al. 2015b), used measures (Videlier et al. 2014; Kelleher et al. 2017) and slightly with age (Urszán et al. 2015b) (see Table 2). The greatest variability was between different arena structures in *Lithobates catesbeianus*, being lowest in the rock-filled enclosure and highest at the plant-filled enclosure (Smith and Doupnik 2005). When tested with different time gaps between repeated measurements, the differences in the repeatability of activity were not apparent (Maes et al. 2012). Furthermore, overall repeatability of activity of larval and post-metamorphic amphibians did not differ much, but no study provided the comparison between multiple life stages of the same individuals, probably due to vast differences in locomotor abilities of larval and post-metamorphic anurans.

### Exploration

Multiple behaviour patterns were observed for exploration. Most of the newts started the trial with a quick escape response and then commenced with the exploration of the outer ring of the arena, rarely visiting the inner parts. A smaller group was startled at first and then explored the inner parts of the arena, eventually reaching the outer ring. The repeatability of exploration in our study was almost identical to the results reported by Gifford et al. (2014) for *Desmognathus brimleyorum*, measured in the same time frame of three weeks. Unfortunately, we were not able to test for the reduction in repeatability with time, as they did. Anuran research showed differences in the consistency of exploration between naïve tadpoles of two different age groups and pretested tadpoles, latter being the highest (Urszán et al. 2015b). The repeatability of post-metamorphic exploration of *Pseudophryne corroboree* (Kelleher et al. 2017) did not differ from larval exploration of *Lithobates catesbeianus* (Carlson and Langkilde 2013) despite different time gaps between repeated measurements. It was also higher than larval exploration of both *Rana temporaria* (Brodin et al. 2013) and *R. dalmatina* (Urszán et al. 2015b), so there does not seem to be any obvious pattern in its strength. The greatest drawback to the comparison of published results is difference in the definition of exploration. It was defined as a buffer around the trajectory of the individual (Brodin et al. 2013), number of visited squares (this study; Carlson and Langkilde 2013; Gifford et al. 2014) or percentage of visited squares (Urszán et al. 2015b). Kelleher et al. (2017) even used three definitions – distance travelled, time spent mobile and number of mobile events, that all showed different repeatability (Table 2). The strength of correlation (repeatability) should therefore not be the main indicator of the suitability of certain measure as the best representative of exploration behaviour. The studies are in need of standardized approaches if they are ever to be compared effectively.

### Boldness

In our study, the repeatability of boldness was present only for the escape response, which is a similar measure to that of Carlson and Langkilde (2013), who counted the number of square sides on the inner portion of the test arena crossed by an individual and which is also a measure of thigmotaxis. Their results are less precise, and the estimated coefficient is lower even though measured with the shorter 24 h breaks between trials (see Table 2). The accuracy of latency of the first movement, as the measure of boldness, was impaired for our study by the inability to consistently raise the glass dome in the centre of the arena at the start of the experiment. We suspect this to be issue in other studies as well, deeming this method impractical. This problem can be mitigated by changing the first movement to a movement longer than one body length (Wilson and Krause 2012), but in our opinion, it is better to choose a completely different option, e.g., shelter use, escape initiation distance or thigmotaxis.

Similar consistency of boldness was reported for *Ambystoma barbouri* and *A. texanum* (Sih et al. 2003) using shelter use as the preferred measure. Looking at other studies, the consistency of boldness varied greatly from 0 to 77 %. The variation was caused by previous experience (Urszán et al. 2015a, 2015b) and the breeding origin (Maes et al. 2012) of the study subjects, and probably by the differences in boldness definition (supported by Kelleher et al. 2017). There does not seem to be any difference between larval and post-metamorphic amphibians or between different time gaps in repeatability measurements (see Table 2).

### Trial repeatability

Activity and exploration significantly decreased with each trial of the experiment, which suggests that habituation may have taken place (but see Carlson and Langkilde 2013). Except for activity, no behavioural trait was repeatable for each trial of the experiment. This could mean that the magnitude of habituation varied individually, i.e., individuality was stronger than habituation (see Fig. 1). For activity, its repeatability for each trial of the experiment was, unexpectedly, followed by low repeatability of both walking and swimming. Other than habituation, another less likely explanation is that the decrease in the expression of behaviour traits could have been caused by insufficient time between the trials of the experiment, allowing newts to remember the last trial. Unfortunately, it was not possible to allow more time between the trials because we feared that newts might switch to the terrestrial phase and change their behaviour. Nevertheless, most amphibian personality studies had an even lower gap between repeated measurements (Table 2). Habituation recovery time is unknown for the studied species. For the common toad, however, Ewert and Kehl (1978) stated that 6–24 h is long enough for recovery from habituation to an artificial rectangular-shaped prey dummy.

**Figure 1.**
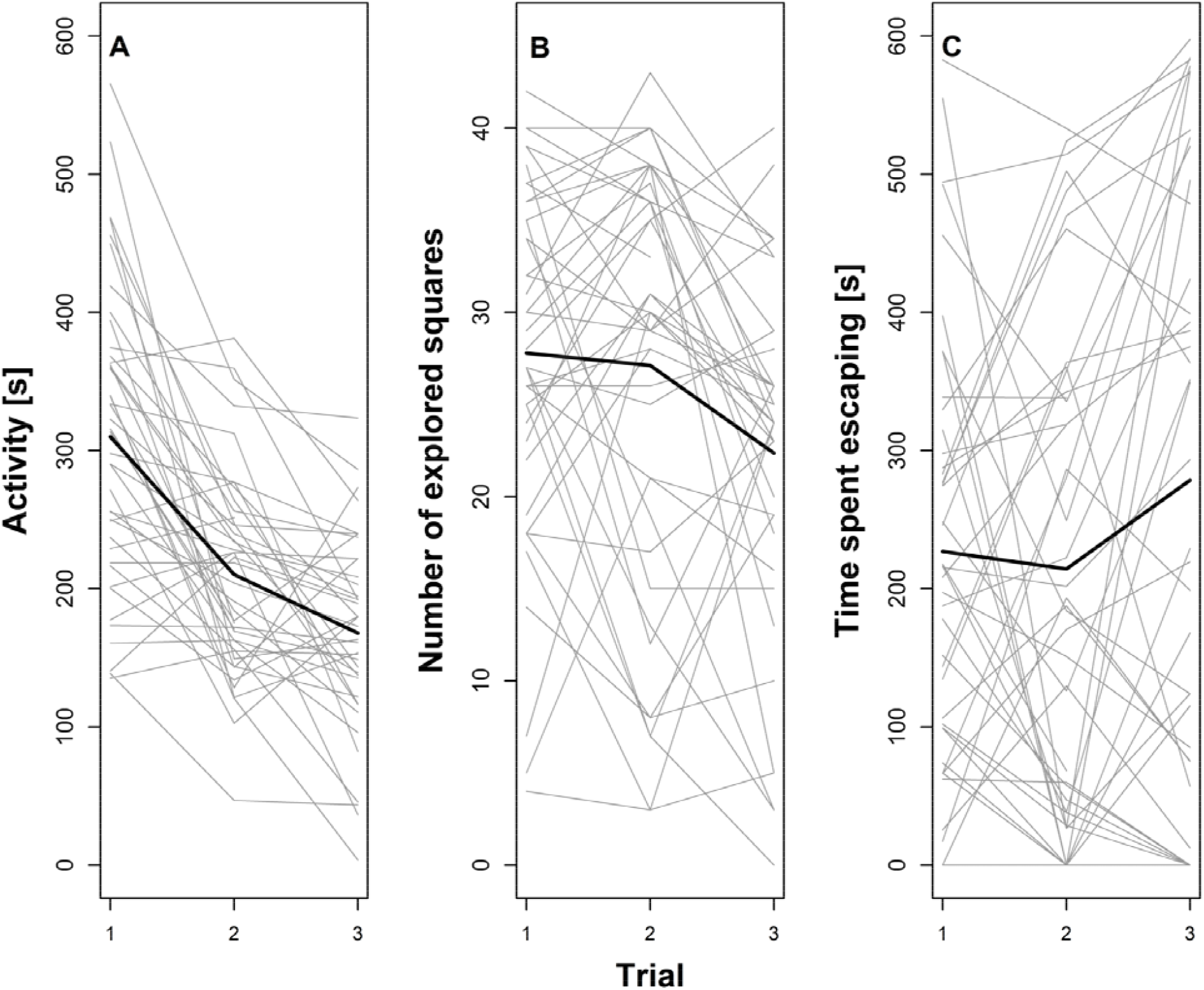
Individuality *vs*. habituation. A – activity between trials, B – exploration between trials, C – escape response between trials.

### Behavioural syndromes

Behavioural syndromes are referred to as suites of correlated behaviours (Sih et al., 2004). According to Kelleher et al. (2018), a total of seven studies assessed behavioural correlations in larval and eight in post-metamorphic amphibians. Activity has been found to correlate with exploration (Koprivnikar et al. 2011; Maes et al. 2012; Wilson and Krause 2012) and boldness (Maes et al. 2012; Wilson and Krause 2012; Urszán et al. 2015a). Boldness has also been found to correlate with exploration (Maes et al. 2012) (but see Brodin et al. 2013), as well as sociability (González-Bernal et al. 2014). Maes et al. (Maes et al. 2012) divided the behaviours using PCA (Principal component analysis) into two axes. The first contained activity and exploration, while the second contained only latency of the first movement. Contrary to their findings, our results show a correlation of all three measured behavioural types (activity, exploration and time spent escaping), potentially creating only one axis (*Kendall’s W* = 0.694, *P* = 0.0001). These behaviours are causally independent and, even being as a part of a single open field test, do not have to be correlated. More movement does not necessarily translate into more squared explored, nor imply more movement near the edge of the arena. In our case, active individuals tended to explore more and were less bold, spending more time escaping. This behaviour might also be a result of a common selective pressure, favouring individuals with a greater insight on the situation in their home pond. The predation rate in the pond was, however, low (pers. obs.), and thus the pressure might be of a reproductive nature. Increase in locomotion activity has been found to benefit in mate searching (Martin et al. 1989), which might result in a correlation with sociability, but we are not aware of any studies researching aggressivity. Because of the absence of behavioural differences between sexes, this pressure might be beneficial for both males and females or at least not harmful for either. Whatever the cause, correlated behaviours, i.e., behaviours that are part of a syndrome should not be studied in isolation because they develop as a group (Sih et al. 2004). To study behaviour syndrome completely, it would also be beneficial to test if the correlations persist in different situations and ecological contexts.

In conclusion, amphibian personality research is sparse, and findings differ considerably in both approach and results. Behavioural consistency is often studied on a small scale in relatively specific conditions, and behavioural correlations are sometimes neglected. We believe that a more complex approach (measuring more types of behaviour) and standardized methodology (i.e., definition of behaviour types, correlation in time and different situations, standard time gap between repeated measurements, number of repeated measurements, duration of experiment and sampling effort, and test arena shape and size) is due. Only then it would be possible to make general assumptions on the global nature and consequences of studied phenomena.

## Acknowledgements

The study was carried out in accordance with permit SZ-092744/2012KUSK/3 issued by the Regional Office of the Central Bohemian Region of the Czech Republic with the support of grants no. 20164234 and no. 20174216, provided by the Internal Grant Agency of the Faculty of Environmental Sciences, Czech University of Life Sciences Prague.

## Author contributions

O. Kopecký captured and cared for newts, and together with P. Chajma participated on the execution of the experiment. P. Chajma analysed the data, prepared all figures and tables and together with J. Vojar wrote the main manuscript. All authors reviewed the manuscript.

